# A Network based Approach to Identify the Genetic Influence Caused by Associated Factors and Disorders for the Parkinsons Disease Progression

**DOI:** 10.1101/482760

**Authors:** Najmus Sakib, Utpala Nanda Chowdhury, M. Babul Islam, Julian M.W. Quinn, Mohammad Ali Moni

## Abstract

Actual causes of Parkinsons disease (PD) are still unknown. In any case, a better comprehension of genetic and ecological influences to the PD and their interaction will assist physicians and patients to evaluate individual hazard for the PD, and definitely, there will be a possibility to find a way to reduce the progression of the PD. We introduced quantitative frameworks to reveal the complex relationship of various biasing genetic factors for the PD. In this study, we analyzed gene expression microarray data from the PD, ageing (AG), severe alcohol consumption (AC), type II diabetes (T2D), high body fat (HBF), hypercholesterolemia (HC), high dietary fat (HDF), red meat dietary (RMD), sedentary lifestyle (SL), smoking (SM), and control datasets. We have developed genetic associations of various factors with the PD based on the neighborhood-based benchmarking and multilayer network topology.

We identified 1343 significantly dysregulated genes in the PD patients compared to the healthy control, where we have 779 genes down regulated and 544 genes up regulated. 69 genes were highly expressed in both for the PD and alcohol consumption whereas the number of shared genes for the PD and the type II diabetes is 51. However, the PD shared 45, 43 and 42 significantly expressed genes with the ageing, high dietary fat and high body fat respectively. The PD shared less than 40 significant transcripts with other factors. Ontological and pathway analyses have identified significant gene ontology and molecular pathways that enhance our understanding of the fundamental molecular procedure of the PD progression. Therapeutic targets of the PD could be developed using these identified target genes, ontologies and pathways. Our formulated methodologies demonstrate a network-based approach to understand the disease mechanism and the causative reason of the PD, and the identification for therapeutic targets of the PD.

## I. Introduction

Parkinson’s disease (PD) is an interminable and gradual degenerative disorder mainly invading the motor system of the central nervous system [1]. It is one of the most common neurodegenerative problems after Alzheimers illness all over the world [2]. The PD is characterized by progressive damage of dopaminergic neurons in the substantia nigra pars compacta and neuronal inclusions composed of -synuclein. These neuronal inclusions which are situated in neuronal perikarya are referred to as Lewy bodies [3]. Subtle early symptoms of the PD appear slowly over time that comprises shaking, rigidity, slowness of movement and walking complications. Patients suffer from complications to walk, talk or even complete other simple daily activities. Sensory, sleep and emotional problems may also evident in the PD. Most often the PD leads to dementia [4]. Approximately 60,000 Americans are diagnosed with the PD each year in the United States which is estimated to reach 1 million by 2020. More than 10 million people are living with the PD worldwide [5]. Although ample discoveries are continuing in this field, the exact causes or risk factors are still poorly understood [2]. Men are more likely to have the PD than women and the domination increases with age. 1% of the total population above 60 years have the PD, whereas only 4% of total cases are estimated at the age of below 50 years [6]. The risk for the PD in humans can significantly be reduced by midlife exercise [7]. Smoking, consumption of alcohol, vitamin D exposure and urate levels are the dominant environmental components that may stimulate the risk of the PD [8]. Moderate doses of caffeine have a protective risk of the PD [9]. Again, the risk of the PD is increased with high total cholesterol at baseline [10]. A diet containing lower saturated fats might lessen the threat of the PD [11]. Type II diabetes weakens the resistance against the PD [12]. The exclusion of dietary red meat boosts the recovery process for several motor functions in the PD affected people [13]. Molecular associations, such as differential gene expressions, protein-protein interactions (PPIs), gene ontologies and metabolic pathways can be genetic associations of various risk factors as the influential causes of the diseases [14], [15]. Any risk factor can be attributed to the disease if they share the common set of differentially expressed genes [16], [17]. But from a proteomics and signaling pathways point of view, they are associated through biological modules such as PPIs, gene ontologies or molecular pathways [18],[19].

Network-based approaches for genetic studies of various diseases have become very popular in recent years [20],[21], [22], [23]. Several genetic studies have been conducted to demonstrate the various risk factors of the PD, but none of them used network-based approaches [24],[25], [26],[27].

In this article, a network based analysis to identify the genetic influence caused by associated factors and disorders for the PD progression is demonstrated utilizing the gene expression profiling, PPI sub-network, gene ontologies and molecular pathways. An extensive study regarding phylogenetic and pathway analysis is also conducted to reveal the genetic associations of the PD.

## II. Materials and Methods

### A. Data

We have analyzed gene expression microarray datasets to identify the association of different factors with the PD at the molecular level. All the datasets used in this study were collected from the National Center for Biotechnology Information (NCBI) Gene Expression Omnibus (https://www.ncbi.nlm.nih.gov/geo/). Ten different datasets with accession numbers: GSE7621, GSE23343, GSE25941, GSE1786, GSE68231, GSE13985, GSE6573, GSE25220, NGSE52553 and GSE4806 are analyzed for this study [28], [29], [30], [31], [32], [33], [34], [35], [36]. The PD dataset (GSE7621) is obtained by RNA extraction and hybridization on Affymetrix microarrays of Substantia nigra tissue from postmortem brain of healthy and the PD patients. The type II diabetes (T2D) dataset (GSE23343) contains gene expression data obtained through extensive analysis after conducting liver biopsies in humans. The hypercholesterolemia (HC) dataset (GSE13985) is obtained from RNA sample of white blood cells of 10 different samples using Affymetrix microarrays. The age (AG) dataset (GSE25941) is a global microarray data from skeletal muscle transcriptome of 28 different subjects. The sedentary lifestyle (SL) dataset (GSE1786) was obtained by expression profiling array from the vastus lateralis muscle using needle biopsies. The high-fat diet (HFD) dataset (GSE68231) is the expression data from human skeletal muscle identifying accumulation of intramyocellular lipid (IMCL). The high body fat (HBF) dataset (GSE6573) is an Affymetrix human gene expression array data from the abdominal fat tissue. The red meat dietary (RMD) intervention dataset (GSE25220) is an Agilent-014850 whole human genome microarray data from human colon biopsies before and after participating in a high red-meat dietary intervention. The alcohol consumption (AC) dataset (GSE52553) is an Affymetrix human gene expression array data of Lymphoblastoid cells from 21 alcoholics and 21 control subjects. The smoking (SM) dataset (GSE4806) is a gene expression profile of T-lymphocytes from smokers and non-smokers.

### B. Method

Nowadays experimental approach based on oligonucleotide microarray data to assess gene expression levels has been found to be an effective and responsive technique to demonstrate the molecular factors of human disorders. In this study, we used this methodology along with global transcriptome analysis to investigate the gene expression profiles of the AD with 8 risk factors and type II diabetes. Since various errors are usually introduced in preparing and analyzing microarray data of different platforms and experimental system, the gene expression data in each sample (disease state or control) need to be normalized. The *Z*-score transform is one of the most widely used normalization methods of gene expression matrix.

Let *x*_*ij*_ be the expression value of *i*-th gene in sample *j*, then normalization using *Z*-score transform is obtained as follows:

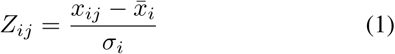

where *σ*_*i*_ and all samples respectively. This transform allows for the direct comparison of gene expression values over different samples and diseases.

In addition to the above *Z*-score transform, we performed linear regression method on the time series data to obtain a joint t-test statistic between two conditions. Data were transformed using *log*_2_ and the linear regression model for calculating the expression level of each gene was as follows:

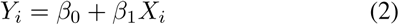

where *Y*_*i*_ is the gene expression value and *X*_*i*_ is a disease state (disease or control). The model parameters *β*_0_ and *β*_1_ were computed using least squares.

To identify differentially expressed genes for both disease and control states we applied unpaired t-test, and significant genes were selected by setting thresholds for p-value to at most 0.05 and absolute log Fold Change (*log*_2_*FC*) value to at least 1.0. The microarray gene expression datasets were collected from NCBI-GEO dataset [36]. All the datasets were analyzed and compared with the normal subject using NCBIs GEO2R online tool to identify corresponding differentially expressed genes.

The web-based visualization software STRING [37] was used for the construction and analysis of the Protein-Protein Interaction (PPI) network which was further analyzed by Cytoscape (v3.5.1) [38]. An undirected graph representation was used for the PPI network, where the proteins were represented by the nodes and the edges symbolized the interactions between the proteins. We performed a topological analysis using Cyto-Hubba plugin [39] to identify highly connected proteins (i.e., hub proteins) in the network and the degree metrics were employed [40]. For further introspection into the metabolic pathways of the PD, we incorporated the pathway and gene ontology analysis on all the differentially expressed genes that were common among the PD and other risk factors and diseases using the web-based gene set enrichment analysis tool EnrichR [41]. In this analysis, the Gene Ontology (GO) Biological Process (BP) and KEGG pathway databases were selected as annotation sources. For statistical significance, the highest adjusted p-value was considered 0.05 to obtain enrichment results. Obtained GO and pathway were further analyzed by Cytoscape.

## III. Results

### A. Gene Expression Analysis

To identify the dysregulated genes due to the PD, the gene expression patterns from substantia nigra tissues of 16 PD patients were analyzed and compared with 9 normal subjects (https://www.ncbi.nlm.nih.gov/geo/geo2r/?acc=GSE7621) [28]. A total of 1343 genes with p-value less than 0.05 and log2 *FC* value greater than 1.0 were found to be differentially expressed compared to healthy subjects where 544 genes were expressively up-regulated and 779 genes were down-regulated.

In order to investigate the association of the PD with 8 risk factors and type II diabetes, we analyzed messenger RNA (mRNA) microarray data separately for all risk factors and diseases. In this analysis, several steps of statistical method have been followed, and the most significant up- and down-regulated genes for each risk factor and disease were selected. We identified a number of differentially expressed genes (1438 in T2D, 958 in ageing, 800 in SL, 739 in HFD, 57 in HC, 824 in HBF, 482 in RMD, 1405 in AC, and 400 in SM) in our analysis.

The common over and under expressed genes among the PD, risk factors and diseases are also detected through a cross-comparative analysis. The findings demonstrated that the PD shares total 51, 45, 26, 43, 4, 42, 20, 69 and 17 prominent genes with T2D, AG, SL, HFD, HC, HBF, RMD, AC, and SM respectively. Two diseasome association networks centered on the PD were built using Cytoscape (v3.5.1) to identify statistically significant associations among these risk factors and diseases [38]. The network shown in Fig. 1 interprets the association among up-regulated and down-regulated genes.

**Fig. 1:**
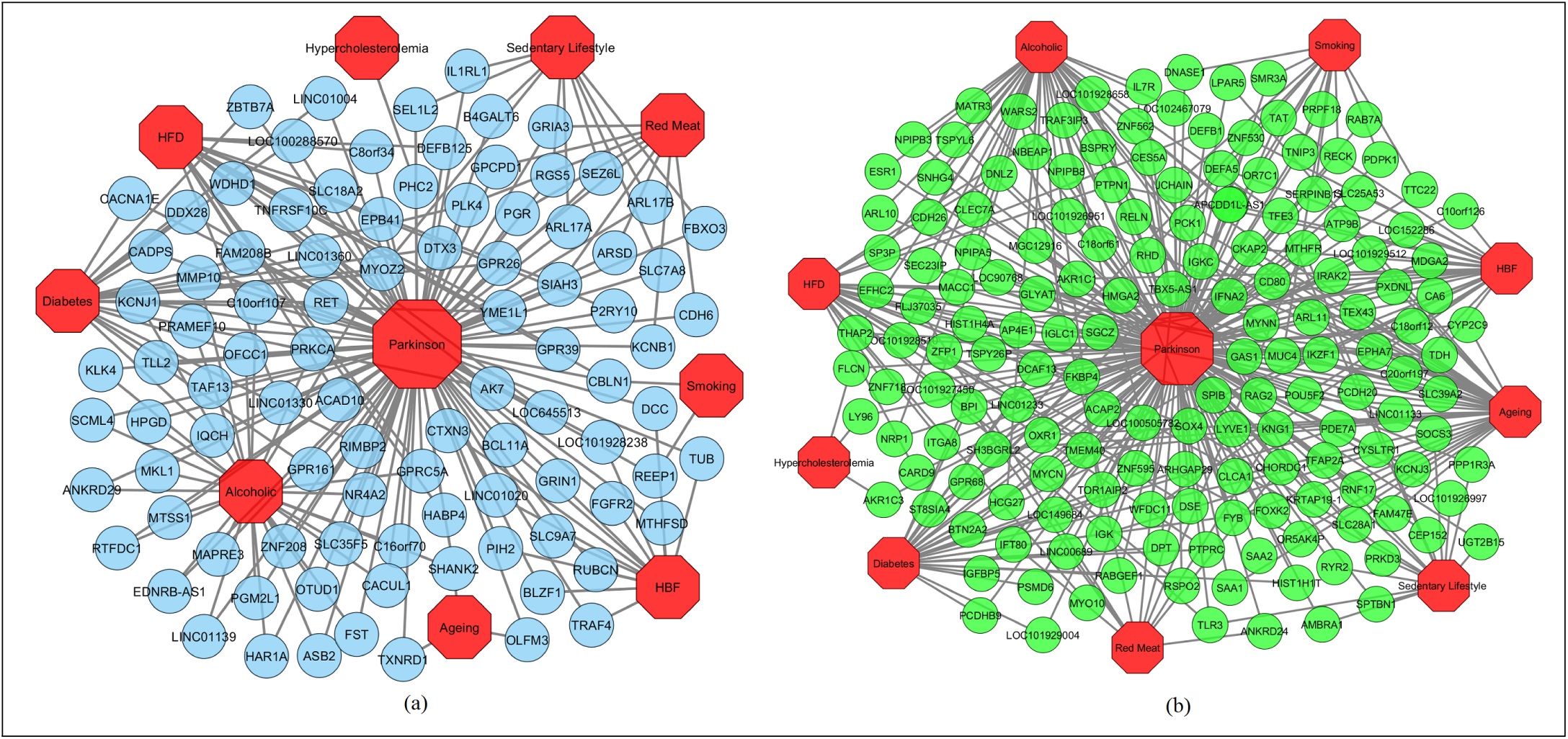
Diseasome network of the Parkinsons disease (PD) with T2D, AG, SL, HFD, HBF, RMD, AC, HC and SM. (a) Red-colored octagon-shaped nodes represent factors and/or disease, and round-shaped sky blue-colored nodes represent up-regulated genes that are common for the PD with the other risk factors and/or diseases. (b) Red-colored octagon-shaped nodes represent factors and/or diseases, and round-shaped green-colored nodes represent down-regulated genes that are common for the PD with the other risk factors and/or diseases. A link is placed between a risk factor or disease and gene if variation of that gene leads to the specific disorder.

In our study, we found that 2 significant genes AK7 and YME1L1 are commonly up-regulated among the PD, T2D and HFD; 2 up-regulated genes GPR26 and PLK4 are common among the PD, T2D, and RMD; 2 significant genes, MYOZ2 and OFCC1 are commonly up-regulated among the PD, T2D and AC. On the other hand, 3-down regulated genes IGK, IGKC and IGLC1 are common among the PD, T2D, AG and AC; 2 down-regulated genes EPHA7 and HMGA2 are common in the PD, AG, HBF and HFD; 3 down-regulated genes APCDD1L-AS1, MUC4 and IGLC1 are common among the PD, AC and HBF; 3 down-regulated genes DCAF13, FKBP4 and PTPRC are common among the PD, T2D, and HFD; 4 down-regulated genes PCK1, KRTAP19-1, FYB and IGLC1 are common among the PD, AG and AC; 3 down-regulated genes CKAP2, EPHA7 and HMGA2 are common among the PD, AG and SL; 4 down-regulated genes SGCZ, IKZF1, MYCN and IGLC1 are common among PD, T2D and HBF; 3 down-regulated genes SOX4, TLR3 and HIST1H4A are common among the PD, HFD and SL; 2 down-regulated genes JCHAIN and IGKC are common among the PD, AG and SM; 2 down-regulated genes ARHGAP29 and BPI are common among the PD, T2D and RMD; 2 down-regulated genes TFE3 and KNG1 are common among the PD, AC and SL; 2 down-regulated genes RHD and IGLC1 are common among the PD, AC and SM; 2 down-regulated genes PTPN1 and HIST1H4A are common among PD, HBF and HFD; 2 down-regulated genes KNG1 and MTHFR are common among the PD, RMD and SL.

### B. Protein-Protein Interaction Network Analysis

The PPI network was constructed using all the distinct 260 differentially expressed genes that were common among the PD, other risk factors and T2D (Fig. 2). Each node in the network represents a protein and an edge indicates the interaction between two proteins. The network is also grouped into 9 clusters representing risk factors and diseases to depict the protein belongings. Notably, each of HIST1H4A and KNG1 proteins belong to the maximum three clusters indicating that they are the common among the PD and other 3 risk factors and T2D, and they interact with other proteins from different clusters. However, each of the proteins MTHFR, GPR26, FKBP4, PTPRC, YME1L1, FYB, IKZF1, PCK1, MYCN, OFCC1, PTPN1 and TLR3 belong to two clusters and interact with other proteins in the network. For topological analysis, a simplified PPI network was constructed using Cyto-Hubba plugin [39] to show 10 most significant hub proteins (Figure-3), which are ESR1, PTPRC, CD80, KNG1, SAA1, GPR68, PRKCA, RET, TLR3 and IL7R. These hub proteins could be the targeted proteins for the drug development.

**Fig. 2:**
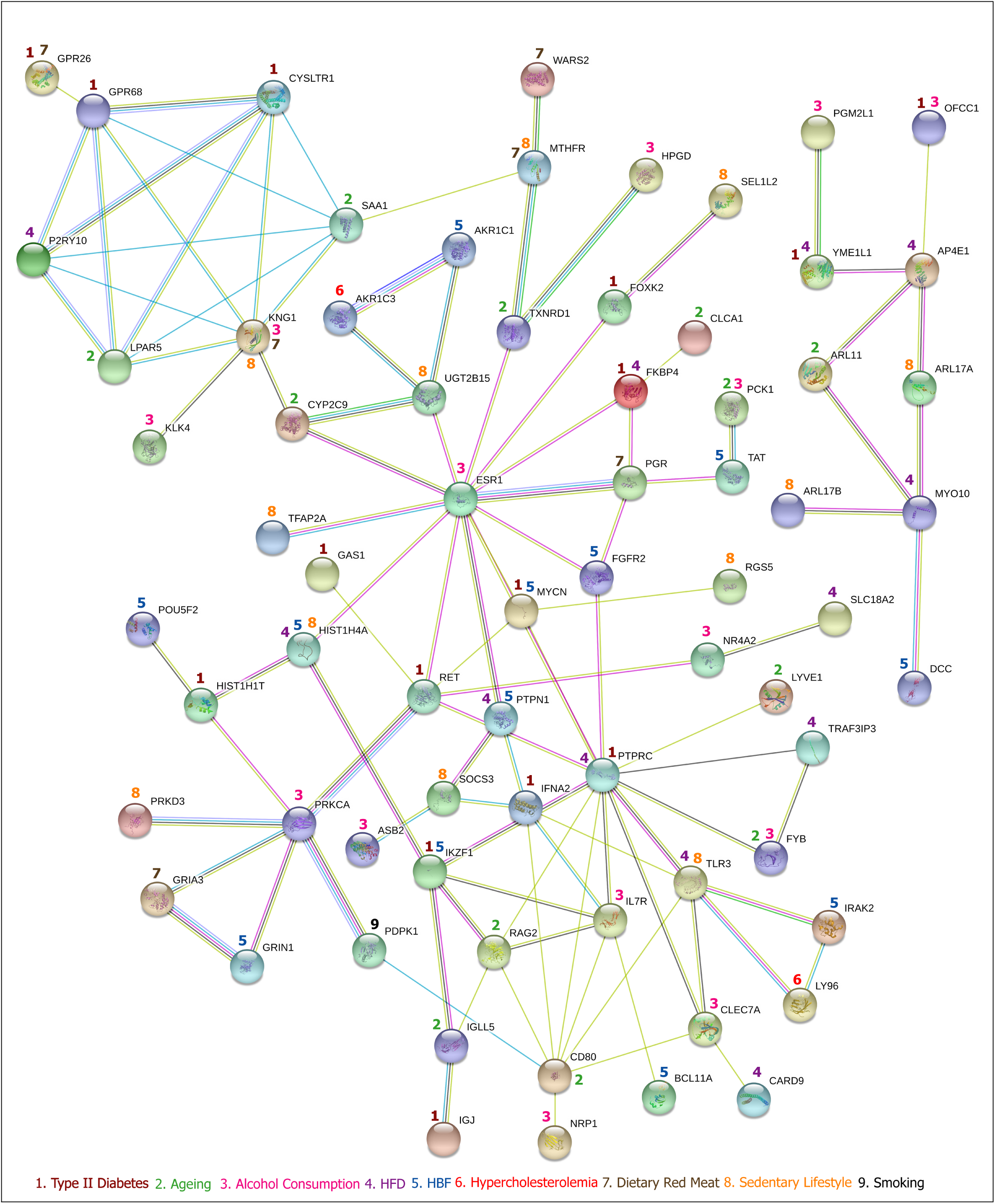
Protein-protein interaction network of commonly dysregulated genes among the PD, 8 risk factors and type II diabetes. Each cluster indicates the gene belongings.

**Fig. 3:**
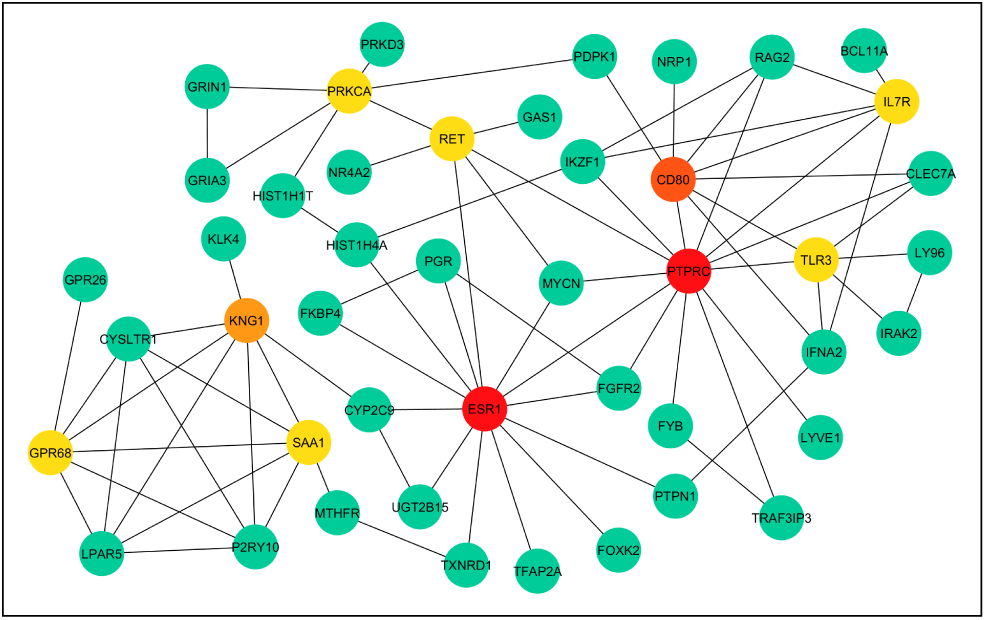
The simplified PPI network of the commonly dysreg-ulated genes between the PD and other risk factors and T2D. The 10 most significant hub proteins are marked as red, orange and yellow.

### C. Pathway and Functional Correlation Analysis

In order to identify the molecular pathways associated with the PD and we have performed pathway analysis on all the differentially expressed genes that were common among the PD, other risk factors and diseases using the KEGG pathway database (http://www.genome.jp/kegg/pathway.html) and the web-based gene set enrichment analysis tool EnrichR [41]. Total 50 pathways were found to be overrepresented among several groups. Notably, two significant pathways that are related to the nervous system have been found which are Glutamatergic synapse (hsa04724) and Serotonergic synapse (hsa04726). These pathways along with some other common pathways are shown in Table I.

**TABLE I:**
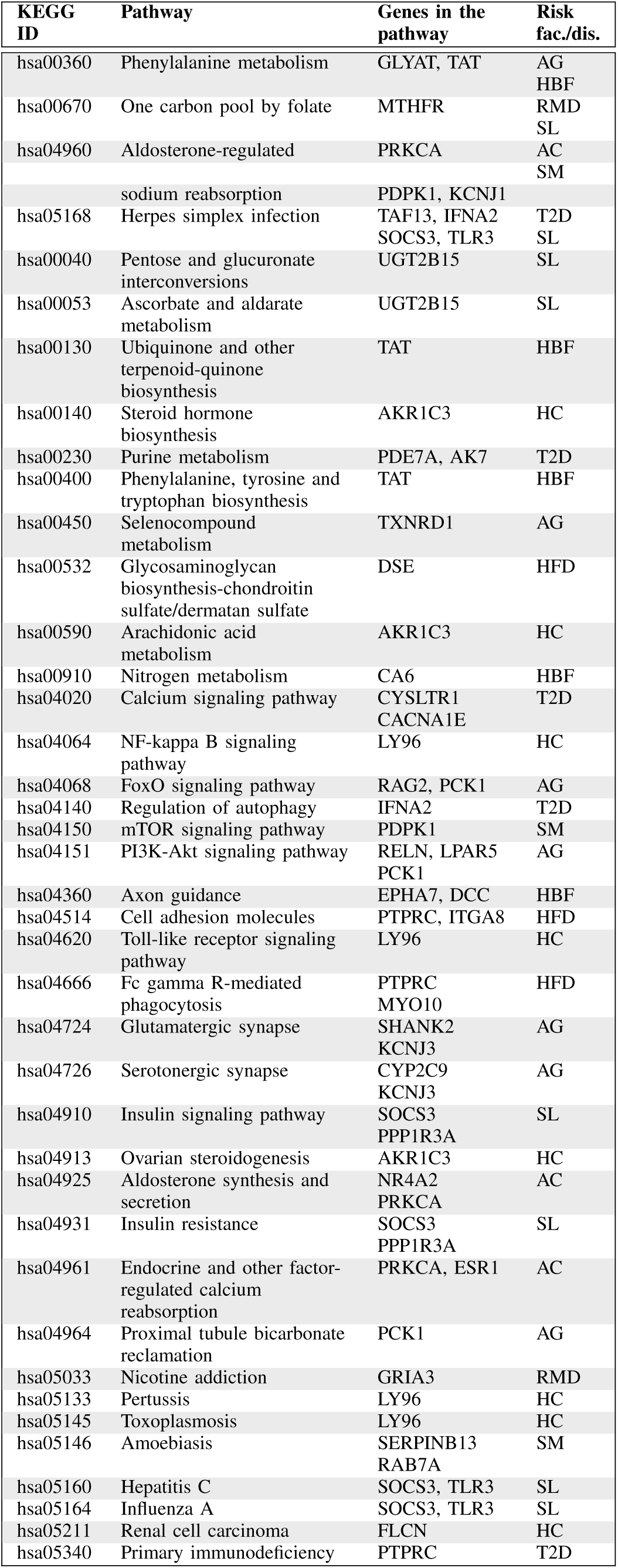
Significant KEGG Pathways related to the Nervous System and Common between the PD and Other Risk Factors and T2D.

Besides, we dug up the overrepresented ontological groups by performing gene biological process ontology enrichment analysis using EnrichR on the commonly dysregulated genes among the PD, other risk factors and diseases. Total 1202 significant gene ontology groups including peripheral nervous system neuron development (GO:0048935), neurotransmitter transport (GO:0006836), neuromuscular synaptic transmission (GO:0007274), peripheral nervous system development (GO:0007422), negative regulation of neurological system process (GO:0031645), regulation of neurotransmitter secretion (GO:0046928), regulation of neuronal synaptic plasticity (GO:0048168), autonomic nervous system development (GO:0048483), sympathetic nervous system development (GO:0048485), neuromuscular process controlling balance (GO:0050885), neuron apoptotic process (GO:0051402), regulation of neurotransmitter transport (GO:0051588) and neuroepithelial cell differentiation (GO:0060563), were observed. Table II summarizes some common genes ontology groups.

**TABLE II:**
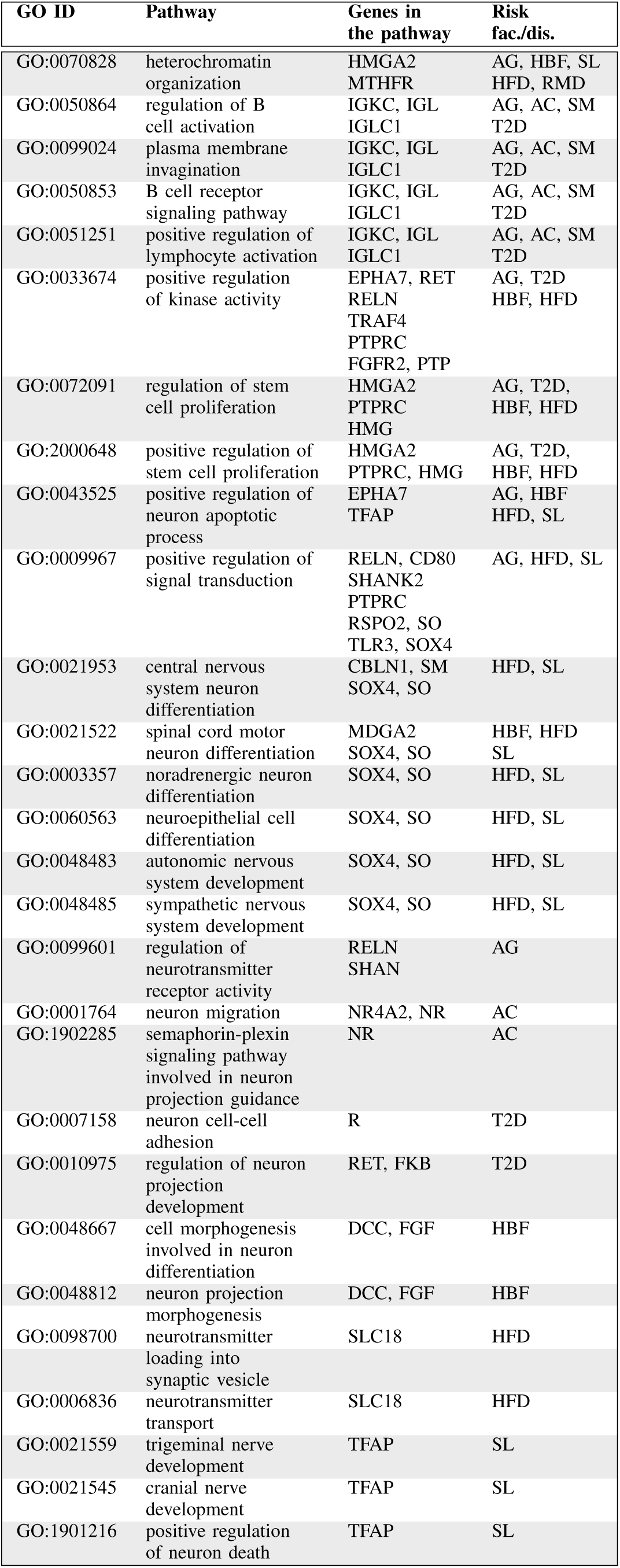
Significant GOs Related to the Nervous System and Common between the PD and Other Risk Factors and T2D.

## IV. Discussion

In this study, we investigated the molecular mechanism of the PD and its genetic association with other risk factors and diseases. For this purpose, we conducted analysis of dysregulation in gene expression of the PD patients, molecular key pathways, ontologies and PPIs. These analyses through network-based approach can unfold novel relationships between the PD and other susceptibility/risk factor. Findings could be very potential and have not been possessed by any previous individual studies. Our outcomes identified several significant genes that yield an opportunity to identify therapeutic targets for the PD. Besides this, our analysis also identified and characterized various biological functions related with these genes.

Our gene expression analysis showed that the PD is strongly associated with alcohol consumption (69), type II diabetes (51 genes), ageing (45 genes), HFD (43 genes) and HBF (42) as they share the maximum number of genes. We constructed and analyzed the PPI network to have a better understanding of the central mechanism behind the PD. For this reason, to construct a PPI network around the differentially expressed genes for our study, we have combined the results of statistical analyses with the protein interactome network. For finding central proteins (i.e., hubs), topological analysis strategies were employed. These identified Hubs proteins might be considered as candidate biomarkers or potential drug targets. From the PPI network analysis, it is observed that 10 hub genes (ESR1, PTPRC, CD80, KNG1, SAA1, GPR68, PRKCA, RET, TLR3 and IL7R) are involved in the PD.

Furthermore, disease related genes play a vital role in the human interactomes via the pathways. In this study, we identified two significant pathways that are associated with the nervous system which are Glutamatergic synapse and Serotonergic synapse. Our study also identified several gene ontologies groups including peripheral nervous system neuron development, neurotransmitter transport, neuromuscular synaptic transmission, peripheral nervous system development, negative regulation of neurological system process, regulation of neurotransmitter secretion, regulation of neuronal synaptic plasticity, autonomic nervous system development, sympathetic nervous system development, neuromuscular process controlling balance, neuron apoptotic process, regulation of neurotransmitter transport and neuroepithelial cell differentiation, which are closely related to the nervous system.

We have also analyzed the differentially expressed genes of each risk factor and disease with Online Mendelian Inheritance in Man (OMIM) and dbGAP databases to validate our identified result using the valid gold benchmark drug-gene associations. Table III summarizes the results. The results corroborate that, the differentially expressed genes of 8 risk factors and type II diabetes cause the PD. As a whole, our findings compensate a major gap of about the PD biology. It will also open up an entry point to establish a mechanistic link among the PD, various risk factors and diseases.

**TABLE III:** Gene-Disease Association Analysis of Differen-tially Expressed Genes of 8 Risk Factors and T2D Using OMIM Database.

| Risk fac./dis. | Gene | Adj. p-value |
| --- | --- | --- |
| AG | UNC13C, RBFOX3, BANK1, STAP1, PLAG1, FAM47E, SNX10, KBTBD11, SNCA, HMGA2 | 1.68E-01 |
| AC | SEMA5A, NEGR1, RAB25, SH3GL2, ADCY2, BRINP1, PLCB4, SEMA6D, MGAT5, ADCY1, STAP1, CTNNA3, MACROD2, MAPT, IGF1R, FGF12, DLG2, NR4A2, GNAQ, GNAS, ARNT | 3.80E-02 |
| T2D | GABBR2, USP36, AKAP13, SEMA6D, CYB5RL, DSG3, RREB1, SGCZ, CYP17A1, HNF4A, HMGA2, BUB1, IGF1R | 5.49E-01 |
| HBF | PTGIR, THRA, HMGA2, ADCY2, DRD1, BUB1, FRMPD4, ZNF646, CTNNA3, MACROD2, TNPO2, SGCZ | 2.03E-01 |
| HC | IQGAP1 | 2.32E-01 |
| HFD | AKAP13, IL1RL2, ITGA8, RREB1, TNPO2, GABRG3, CREBBP, HNF4A, HMGA2 | 3.47E-01 |
| RMD | C9, ULK2, ZNF385B, GABRG3, SNCA, HMGA2, ADCY2 | 2.03E-01 |
| SL | GAK, NELL1, BRINP1, COL4A1, VWA5A, PLEKHM1, PHACTR2, TFAP2A, HSP90AA1, CREB1, THRA, TSHR | 2.44E-01 |
| SM | NR4A2, OPRD1, SMAD1, BUB1, PAK1, CREM, TNPO2 | 1.15E-01 |

## V. Conclusion

In this study, genomic data is considered to identify the genetic association of various diseasome relationships with the PD. Our findings elicit that the network methods can illustrate disease progression that yields a potential advancement towards better insight of the origin and evolution of the PD. Detecting the complex relationship of various risk factors with the disease may expose new and useful information important for having a better understanding of overall mechanism as well as planning remedy strategy of the PD. Making genomic based recommendations for accurate disease diagnosis and effective treatment can be boosted by the approaches demonstrated in this study. This enhancement may eventually lead genomic information-based personalized medicine for the more precise insight into disease mechanism, and thus an opportunity to determine accurate diagnosis, treatment and remedy from the PD disease.

## References

[1] National Institute of Neurological Disorders, and Stroke (US). Parkinson’s disease: Hope through research. No. 94–139. National Institute of Neurological Disorders and Stroke, National Institutes of Health, 1994.

[2] Kalia LV, Lang AE (August 2015), “Parkinson’s disease". Lancet. 386 (9996):896–912.

[3] Dickson, Dennis W, “Parkinson’s disease and parkinsonism: neuropathology,” Cold Spring Harbor perspectives in medicine (2012):a009258.

[4] Sveinbjornsdottir, Sigurlaug, “The clinical symptoms of Parkinson’s disease,” Journal of neurochemistry 139 (2016):318–324.

[5] http://www.parkinson.org/understanding-parkinsons/causes-and-statistics

[6] Tysnes, Ole-Bjorn, and Anette Storstein, “Epidemiology of Parkinson’s disease,” Journal of Neural Transmission 124.8 (2017):901–905.

[7] Ahlskog, J. Eric, “Does vigorous exercise have a neuroprotective effect in Parkinson disease?” Neurology 77.3 (2011):288–294.

[8] Kalia LV, Lang AE (2015) Parkinson’s disease. Lancet 86(9996):896–912.

[9] Ascherio, Alberto, et al, “Prospective study of caffeine consumption and risk of Parkinson’s disease in men and women” Annals of Neurology: Official Journal of the American Neurological Association and the Child Neurology Society50.1 (2001):56–63.

[10] Hu, G., et al., “Total cholesterol and the risk of Parkinson disease,” Neurology 70.21 (2008):1972–1979.

[11] Kamel, Freya, et al., “Dietary fat intake, pesticide use, and Parkinson’s disease,” Parkinsonism & related disorders 20.1 (2014):82–87.

[12] Hu, Gang, et al., “ype 2 diabetes and the risk of Parkinson’s disease,” Diabetes care (2007).

[13] Coimbra, Cicero Galli, and Virginia Berlanga Campos Junqueira, “High doses of riboflavin and the elimination of dietary red meat promote the recovery of some motor functions in Parkinsons disease patients,” Brazilian Journal of Medical and Biological Research 36.10 (2003):1409–1417.

[14] Rzhetsky, Andrey, et al., “Probing genetic overlap among complex human phenotypes,” Proceedings of the National Academy of Sciences 104.28 (2007.):11694–11699.

[15] Lee, D-S., et al., “The implications of human metabolic network topology for disease comorbidity,” Proceedings of the National Academy of Sciences (2008.).

[16] Goh, Kwang-Il, et al., “he human disease network,” Proceedings of the National Academy of Sciences 104.21 (2007.):8685–8690.

[17] Feldman, Igor, Andrey Rzhetsky, and Dennis Vitkup, “Network properties of genes harboring inherited disease mutations,” Proceedings of the National Academy of Sciences 105.11 (2008.):4323–4328.

[18] Lage, Kasper, et al, “A human phenome-interactome network of protein complexes implicated in genetic disorders,” Nature biotechnology 25.3 (2007.):309.

[19] Suthram, Silpa, et al., “Network-based elucidation of human disease similarities reveals common functional modules enriched for pluripotent drug targets,” PloS computational biology 6.2 (2010):e1000662.

[20] Moni, Mohammad Ali, and Pietro Lio’, “Network-based analysis of comorbidities risk during an infection: SARS and HIV case studies,” BMC bioinformatics 15.1 (2014.):333.

[21] Moni, Mohammad Ali, and Pietro Lio, “Genetic profiling and comorbidities of Zika infection,” The Journal of infectious diseases 216.6 (2017):703–712.

[22] orkamani, Ali, Eric J. Topol, and Nicholas J. Schork, “Pathway analysis of seven common diseases assessed by genome-wide association,” Genomics 92.5 (2008.):265–272.

[23] Baranzini, Sergio E., et al., “Pathway and network-based analysis of genome-wide association studies in multiple sclerosis,” Human molecular genetics 18.11 (2009.):2078–2090.

[24] International Parkinson Disease Genomics Consortium, “Imputation of sequence variants for identification of genetic risks for Parkinson’s disease: a meta-analysis of genome-wide association studies,” The Lancet 377.9766 (2011.):641–649.

[25] Klein, Christine, and Andreas Ziegler, “From GWAS to clinical utility in Parkinsons disease,” The Lancet 377.9766 (2011.):613–614.

[26] Hardy, John, “Genetic analysis of pathways to Parkinson disease,” Neuron 68.2 (2010.):201–206.

[27] Brs, Jos, Rita Guerreiro, and John Hardy, “SnapShot: genetics of Parkinson’s disease,” Cell 160.3 (2015.):570–570.

[28] Lesnick TG, Papapetropoulos S, Mash DC, Ffrench-Mullen J et al., “A genomic pathway approach to a complex disease: axon guidance and Parkinson disease,” PloS Genet 2007. Jun; 3(6):e98.

[29] Misu H, Takamura T, Takayama H, Hayashi H et al., “A liver-derived secretory protein, selenoprotein P, causes insulin resistance,” Cell Metab 2010. Nov 3;12(5):483–95.

[30] Raue U, Trappe TA, Estrem ST, Qian HR et al., “Transcriptome signature of resistance exercise adaptations: mixed muscle and fiber type specific profiles in young and old adults,” J Appl Physiol (1985) 2012 May;112(10):1625–36.

[31] Radom-Aizik S, Hayek S, Shahar I, Rechavi G et al., “Effects of aerobic training on gene expression in skeletal muscle of elderly men,” Med Sci Sports Exerc 2005 Oct;37(10):1680–96.

[32] Kakehi S, Tamura Y, Takeno K, Sakurai Y et al., “Increased intramyocellular lipid/impaired insulin sensitivity is associated with altered lipid metabolic genes in muscle of high responders to a high-fat diet,” Am. J. Physiol Endocrinol Metab2016 Jan 1; 310(1):E32–40.

[33] Herse F, Dechend R, Harsem NK, Wallukat G et al., “Dysregulation of the circulating and tissue-based renin-angiotensin system in preeclampsia,” Hypertension 2007. Mar; 49(3):604–11.

[34] Hebels DG, Sveje KM, de Kok MC, van Herwijnen MH et al., “N-nitroso compound exposure-associated transcriptomic profiles are indicative of an increased risk for colorectal cancer,” Cancer Lett 2011. Oct 1; 309(1):1–10.

[35] McClintick JN, Brooks AI, Deng L, Liang L et al., “Ethanol treatment of lymphoblastoid cell lines from alcoholics and non-alcoholics causes many subtle changes in gene expression,” Alcohol 2014. Sep; 48(6):603–10.

[36] Bttner P, Mosig S, Funke H, “Gene expression profiles of T lymphocytes are sensitive to the influence of heavy smoking: A pilot study,” Immunogenetics2007. Jan; 59(1):37–43.

[37] Szklarczyk, Damian, et al., “The STRING database in 2017: quality-controlled proteinprotein association networks, made broadly accessible,” Nucleic acids research (2016): gkw937.

[38] Smoot, Michael E., et al., “Cytoscape 2.8: new features for data integration and network visualization,” Bioinformatics27.3 (2010.): 431–432.

[39] Chen, Shu-Hwa, et al., “cyto-Hubba: A Cytoscape plug-in for hub object analysis in network biology,” 20th International Conference on Genome Informatics, 2009

[40] Calimlioglu, Beste, et al., “Tissue-specific molecular biomarker signatures of type 2 diabetes: an integrative analysis of transcriptomics and proteinprotein interaction data,” Omics: a journal of integrative biology 19.9 (2015.): 563–573.

[41] Kuleshov, Maxim V., et al., “Enrichr: a comprehensive gene set enrichment analysis web server 2016 update,” Nucleic acids research 44.W1 (2016): W90–W97.

